# The maintenance of self-incompatibility and the genetic architecture of inbreeding depression

**DOI:** 10.1101/2022.03.24.485676

**Authors:** Daniel J. Schoen, Sarah J. Baldwin

## Abstract

- Inbreeding depression plays a fundamental role in evolution. To help detect and characterize viability loci that underlie inbreeding depression, we forced self-pollinated plants from self-incompatible populations of *Leavenworthia alabamica* to produce families of progeny that were genotyped at hundreds of mapped single nucleotide polymorphism (SNP) loci.
- Bayesian analysis of segregation data for each SNP was used to explore support for different dominance and selection coefficients at linked viability loci in different genomic regions.
- There was some support for overdominance (or pseudo-overdominance) at a few viability loci, as well as for recessiveness and underdominance. One recessive viability locus mapped to the genomic region of the novel self-incompatibility locus in *Leavenworthia alabamica*, but in general there was no support for strongly recessive viability loci of major effect.
- The results are consistent with earlier findings showing that inbreeding depression is recalcitrant to purging in *Leavenworthia alabamica*. The results also help account for the maintenance of self-incompatibility in this species and are consistent with expectations from evolutionary genetic theory that recessive, deleterious alleles linked to loci under balancing selection are sheltered from selection.

## Introduction

Inbreeding depression is the reduction of fitness in progeny produced by consanguineous mating versus mating between unrelated parents (Wright, 1977). Inbreeding depression plays a fundamental role in crop production, conservation of natural populations, and mating system evolution (Charlesworth & Willis, 2009; Frankham, 2010; Lloyd, 1979; Lande & Schemske, 1985). While recessive deleterious mutations are thought to play a major role in inbreeding depression, overdominance has also been implicated (Wright, 1977; Charlesworth & Willis, 2009).

Lande & Schemske (1985) considered the evolutionary consequences of inbreeding depression and the mating system. Their theoretical work predicted when there is selective purging of recessive lethal and near-lethal alleles, populations of self-fertilizing organisms should exhibit reduced equilibrium levels of inbreeding depression. Charlesworth & Charlesworth (1987) extended the study of purging of inbreeding depression to many deleterious alleles of milder effect with a broader range of dominance and found that fairly high levels of inbreeding depression can be maintained in populations of self-fertilizing organisms. For species with self-incompatibility (SI), theory predicts that an additional source of inbreeding depression is due to the linkage of deleterious recessive alleles to self-recognition loci (Uyenoyama, 1997). In gametic self-incompatibility systems the self-recognition locus (S-locus) region is maintained as heterozygous, thereby hindering the selective purging of recessive deleterious mutations, and leading to “sheltered genetic load” in this region. Likewise, in sporophytic self-incompatibility systems, depending on the dominance levels of the S-allele, the S-locus region is expected to be maintained as heterozygous and harbour sheltered S-linked load (Schierup *et al*., 1997; Llaurens *et al*., 2009). Theory has shown that sheltered load may affect the trajectory of mating system evolution (Porcher & Lande, 2005; Gervais *et al*., 2014).

Few studies have provided information on the number and genomic locations of the viability loci responsible for the expression of inbreeding depression, and there is little information on the selection and dominance coefficients of the viability effects, factors important to predicting the evolutionary trajectory of inbreeding depression. For SI species, there is even less information of this kind (though see Karkkainnen *et al*., 1999).

The genus *Leavenworthia* (Brassicaceae) is a group of self-incompatible and self-compatible species that has received considerable attention by evolutionary biologists interested in plant mating system evolution due to both inter- and intraspecific variation in the mating system (Lloyd, 1965; Busch, 2005; Busch *et al*., 2011; Herman *et al*., 2012; Chantha *et al*., 2013). Self-incompatible species in this genus exhibit significant levels of inbreeding depression (Charlesworth *et al*., 1994; Busch, 2005). Recently, Baldwin & Schoen (2018) conducted experiments in which plants from self-incompatible populations of *L. alabamica* were subjected to several generations of selfing. They found that inbreeding depression expressed in these experiments is resistant to purging.

Marker-based approaches to studying the genetic architecture of inbreeding depression can provide information on the dominance and selective coefficients at individual viability loci (Hedrick & Muona, 1990; Fu & Ritland, 1994a,b), which in turn can help shed light on how inbreeding depression evolves. When available, such information can be coupled with recombination and physical maps to reveal the approximate genomic location of viability loci (Mitchell-Olds, 1995; Remington & O’Malley, 2000; Karkkainnen *et al*., 1999). Here we extract information on the genetic architecture of inbreeding depression in *Leavenworthia alabamica* using a genetic marker-based approach. We discuss the relevance of our results to mating system evolution and to previous findings that have used the marker-based approaches to examine the genetic architecture of inbreeding depression in plants.

## Methods

### Plant growth and production of self-fertilized progeny

Two self-incompatible *Leavenworthia alabamica* plants from separate populations in Alabama were used in this work. The first was from the Hatton population (Latitude, Longitude: 34.509931, −87.442041), while the second was from the M32 population (Latitude, Longitude 34.489356, −87.63559) (Baldwin & Schoen, 2018). These plants were grown in separate 9” pots in the McGill University Phytotron and forced to self-fertilize by bud or salt pollinations as described in Chantha *et al*. (2013). Up to 30 pollinations were carried out per plant two times per week. All seeds were collected at maturity, stratified in a dry environment for at least two months, and germinated. The progeny from these two separate seed families (Families 1 and 2, hereafter) were grown to the flowering stage. Sample sizes were 91 and 92 progeny/family, respectively.

### Genotyping-by-sequencing (GBS)

Young leaf tissue samples from the progeny were collected from flowering stage plants and stored at −80°C. DNA was extracted from the leaf tissue with Qiagen plant mini-DNA kits. Extractions were repeated if concentrations were lower than 20 ng/ul and were diluted to 30-50 ng/ul if over 50 ng/ul.

The genotyping-by-sequencing (GBS) method was used to produce fragments for genotyping with restriction-site associated polymorphisms (Elshire *et al*., 2011) using the protocol of Wallace & Mitchell (2017). The standard protocol was used to generate libraries with 1.44 ng PstI per 100 ng DNA and 15-18 cycles of PCR. Fragments were then sequenced on one lane of an Illumina High-Seq run (Family 1, N = 91 and Family 2, N = 90). FASTQC was used to evaluate the basic quality of the sequencing runs. Sequence reads begin with a barcode on the 5’ end identifying the plant of origin, then the restriction site, the sequence to be used for genotyping, followed by another restriction site, and finally the Illumina universal adapter. Reads were demultiplexed using Process Radtags in STACKS v. 2.59. The *Leavenworthia alabamica* reference assembly version 2.5 (Haudry *et al*., 2013, CoGe platform: Genome ID 20247) was indexed and sorted with Burrows-Wheeler Aligner Tool, BWA-0.7.17 (Li & Durbin, 2010), and reads were aligned to the reference genome with BWA-0.7.17. SNP locus segregation ratios derived from the two families were obtained by producing a catalog of segregating loci using the GSTACKS and POPULATIONS routines in STACKS v. 2.59 (Catchen *et al*., 2013) running in R (version 4.1.1). The in-house scripts and associated information used for processing the data are available in the Supplementary Materials section (Table S1).

### SNP filtering and treatment of potential genotyping errors

Previous genomic analysis of *Leavenworthia alabamica* indicates that this species underwent whole genome triplication during its evolutionary history, and chromosome painting with *Arabidopsis thaliana* BAC probes suggests that that some triplicated genome segments were retained as duplicates or triplicates (Haudry *et al*., 2013). The reference genome assembly in this species is only partially complete (N50 = 309 kb); while the haploid chromosome number of *L. alabamica* is 11, the genome assembly consists of 3,321 scaffolds. It comprises *ca*. 174 million bp in total of the expected 315 million bp genome size, the latter estimated from flow cytometry (Brian Husband, pers. comm.). Over half of the scaffolds are short, with lengths 5,000 bp or smaller, though larger ones are > 1M bp. While there is sequence divergence of the duplicated (or triplicated) regions, longer scaffolds in the reference genome likely represent separate genomic regions (Haudry *et al*., 2013). But many shorter duplicated (or triplicated) regions of the genome with extended identity have likely been collapsed into single (artifactual) contigs. GBS products from separate genomic regions that map to such short, artifactual contigs would be expected to yield heterozygote genotype calls. To minimize these artifacts we: (1) filtered out heterozygotes when allele balance fell below 0.333 or above 0.667 using the R- package SNPfiltr v. 1.00 (DeRaad, 2022), (2) restricted our analysis to SNPs that map to scaffolds > 100 kbp in length (these account for 73% of the *L. alabamica* assembly); (3) filtered out SNP loci that exhibited only heterozygotes when adjacent SNP loci on the same scaffold exhibited segregation ratios that were not significantly different from 1:2:1 by Chi-square tests, on the basis that evidence of strong selection should be shared by adjacent SNP loci; and (4) restricted our analysis to loci where data were available from > 90% of the progeny of a cross since genotyping errors are more likely to have occurred when few progeny per family are genotyped successfully.

### Selection and dominance coefficients at linked viability loci

Linkage groups (each containing a minimum of three SNP loci) were constructed separately for Families 1 and 2 with the software package ‘qtl’ version 1.50 (Broman *et al*., 2003) running in R (version 4.1.1). A viability locus located adjacent to a given SNP locus will likely affect not only the segregation ratio at that SNP but also at other nearby SNPs. The linkage maps allowed us to examine segregation ratios of each SNP in relation to that of other nearby SNPs. To arrive at a conservative estimate of the number of viability loci, we limited our analysis to those SNPs that exhibited departures from the expected (null) 1:2:1 segregation ratio at a false discovery rate (Benjamini & Hochberg, 1995) of *q* = 0.05 and 0.10. For linkage groups where departures from a 1:2:1 ratio were observed, we selected the marker showing the most extreme departure for the analysis of selection and dominance coefficients at the linked viability locus.

We note that the analysis of marker locus segregation ratios to estimate selection and dominance at viability loci linked to marker loci assumes departure from the expected 1:2:1 Mendelian segregation ratio is due to viability selection, rather than to meiotic drive and/or gametic selection. The parent plant that is self-fertilized is assumed to be heterozygous at a selectively neutral marker locus *(M)* and a linked viability locus (*V*)—e.g., the parent has the genotype *M_1_V_1_/M_2_V_2_*, and the genotypes *V_1_/V_1_, V_1_/V_2_*, and *V_2_/V_2_* are associated with the fitness values 1-*s*, *1-hs*, and 1, respectively, where *s* is the selective coefficient or the proportional amount by which selection at this locus reduces viability in the homozygote *V*_2_/*V*_2_ (0 ≤ *s* ≤ 1), and *h* is the dominance coefficient. A value of *h* < 0 implies overdominance, *h* = 0 implies fully recessive, 0 ≤ *h* ≤ 0.5 implies partially recessive, *h* = 0.5 implies additive, 0.5 ≤ *h* ≤ 1.0 implies partial dominance, and *h* > 1 implies underdominance. The marker and viability loci are assumed to be linked with recombination fraction *c*.

A Bayesian statistical genetic approach was developed to explore the posterior support for different values of *h* and *s* at viability loci based on the observed numbers of marker locus genotypes. Fu & Ritland (1994a) showed that following self-fertilization, the post-selection genotype frequencies expected for three marker genotypes (*p_11_* for *M*_1_/*M*_1_, *p*_12_ for *M*_1_/*M*_2_, and *p_22_* for *M*_2_/*M*_2_) are:

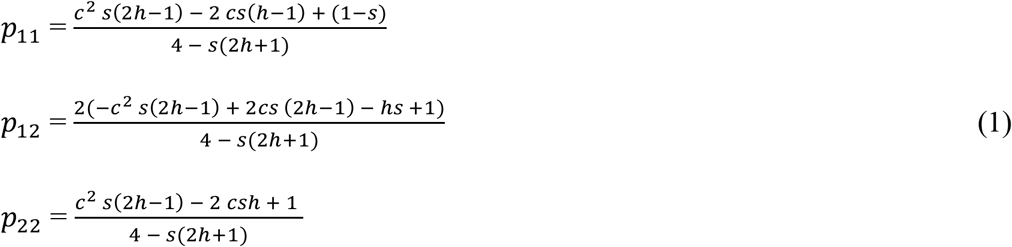

In these equations *p*_11_ denotes the frequency of the least common homozygote genotype. Note that when *s* = 0, the marker genotype frequencies give the null 1:2:1 ratio.

The data are the numbers of each of the three marker genotypes observed among the surviving progeny of the self-fertilized parent (*N*_11_ = number of *M*_1_/*M*_1_, number of observed, *N_12_* = number of *M_1_/M_2_* observed, and *N*_22_ = number of *M_2_/M_2_* observed). The probability (likelihood) given the data can be written:

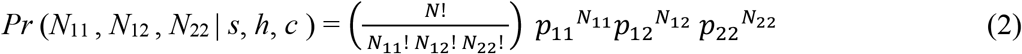

where *N* is the sum of *N*_11_, *N*_12_, and *N*_22_. The joint density and posterior probability are, respectively:

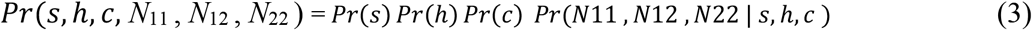

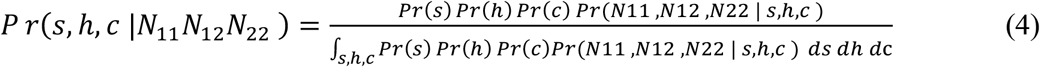

To explore values of the parameters using a Bayesian approach, we chose uniform priors with 0 < *s* < 1, −100 < *h* < 100, and 0 < *c* < 0.2. The broad prior for *h* allows for the possibility of extreme overdominance and underdominance. The prior for *c* (less than the theoretical upper limit of *c* = 0.5) is justified based on the average distance between markers of < 5 cM in this study. A program written in the R statistical language (version 4.1.1) to conduct Markov Chain Monte Carlo method with the Metropolis-Hastings sampler (Weir, 1996) (Supplementary Materials: Note 1). The program was used to “integrate out” the nuisance parameter *c*, allowing the posterior probabilities to be estimated for different values of *s* and *h*. For the required burn-in period, we ran the sampler 5.5 million times before examining the posterior probability distributions. Tracking the parameter values during the runs showed that the sampler had stabilized by the end of this burn-in period.

### The S-locus region of *Leavenworthia alabamica* and locations of viability loci

The *Leavenworthia alabamica* S-locus region has previously been cloned and sequenced from the same plant material that was used to produce the sequence information in the genome assembly in the present work (GenBank accession KC981242.1) (Chantha *et al*., 2013). This allowed us to locate the individual *L. alabamica* genome assembly scaffold containing the S-locus. The BLAST algorithm (Altschul *et al*., 1990) implemented in software package Geneious Prime (version 2.2) was used to conduct the search of *L. alabamica* reference genome version 2.5 using both *L. alabamica* S-locus genes *(Lal2* and *LaSCRL;* Chantha *et al*., 2013). Selection and dominance coefficients were estimated for all potential viability loci linked to SNPs in this region that showed significant departures from 1:2:1 were analysed.

### Statistical power test and different modes of gene action

The analysis of the number and mode of action pf the linked viability loci hinges on the ability to detect departure from the expected 1:2:1 ratio obtained when self-fertilizing a plant with heterozygous marker loci. Given that the sample size of progeny per family in our work was limited to a maximum of 91 progeny (the maximum number we could obtain by forced selfing of each of the SI parent plants used), we were interested in the statistical power of our study to detect selection and characterize the mode of gene action. We did this by producing expected segregation ratios for a range of different values of the parameters in equation (1) followed by standard methods (Cohen, 1988) for examining statistical power, as implemented in the R program package ‘pwr’ version 1.3 (Supplementary Materials: Note 2).

## Results

### SNP segregation ratios, linkage map locations, and significance testing

We found 315 mappable SNP loci in Family 1, and 237 in Family 2. In Family 1 we found 18 linkage groups with 3 or more SNPs, ranging in length from 2 to 201 cM, for a total map length of 901 cM. In Family 2 we found 16 linkage groups with 3 or more SNPs, ranging in length from 3 to 88 cM, for a total map length of 229 cM. SNP grouping by linkage group was consistent in the two families (see Supplementary Tables S2 and S3 for linkage group, scaffold positions, and segregation patterns of SNPs). SNP segregation patterns along with and their - log_10_ significance levels from the Chi-square test of goodness-of-fit to the null 1:2:1 segregation ratio varied within each linkage group are shown in Figures 1 and 2.

**Figure 1.**
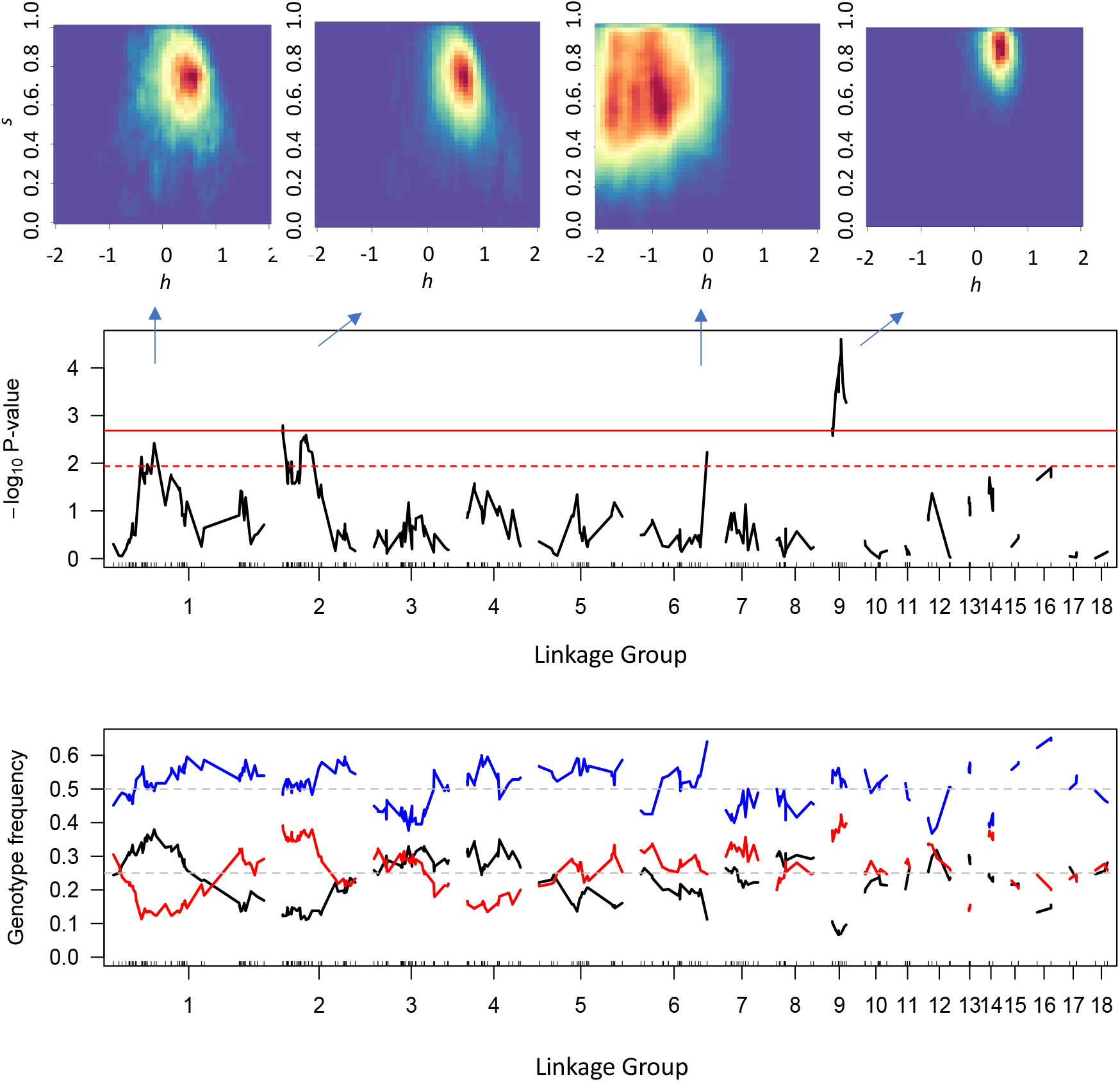
−Log_10_ significance levels from Chi-square goodness-of-fit tests to the 1:2:1 segregation ratio for SNPs in linkage groups of Family 1. Red solid and dashed lines indicate significance levels corresponding to the false discovery rate of *q* = 0.05 and 0.10, respectively. Below are the segregation ratios for the SNPs. Blue denotes heterozygote, red and black denote homozygotes. Above are heat maps of posterior probability densities for dominance (*h*) and selection (*s*) coefficients of inferred viability loci in linkage groups (positions indicated by arrows).

**Figure 2.**
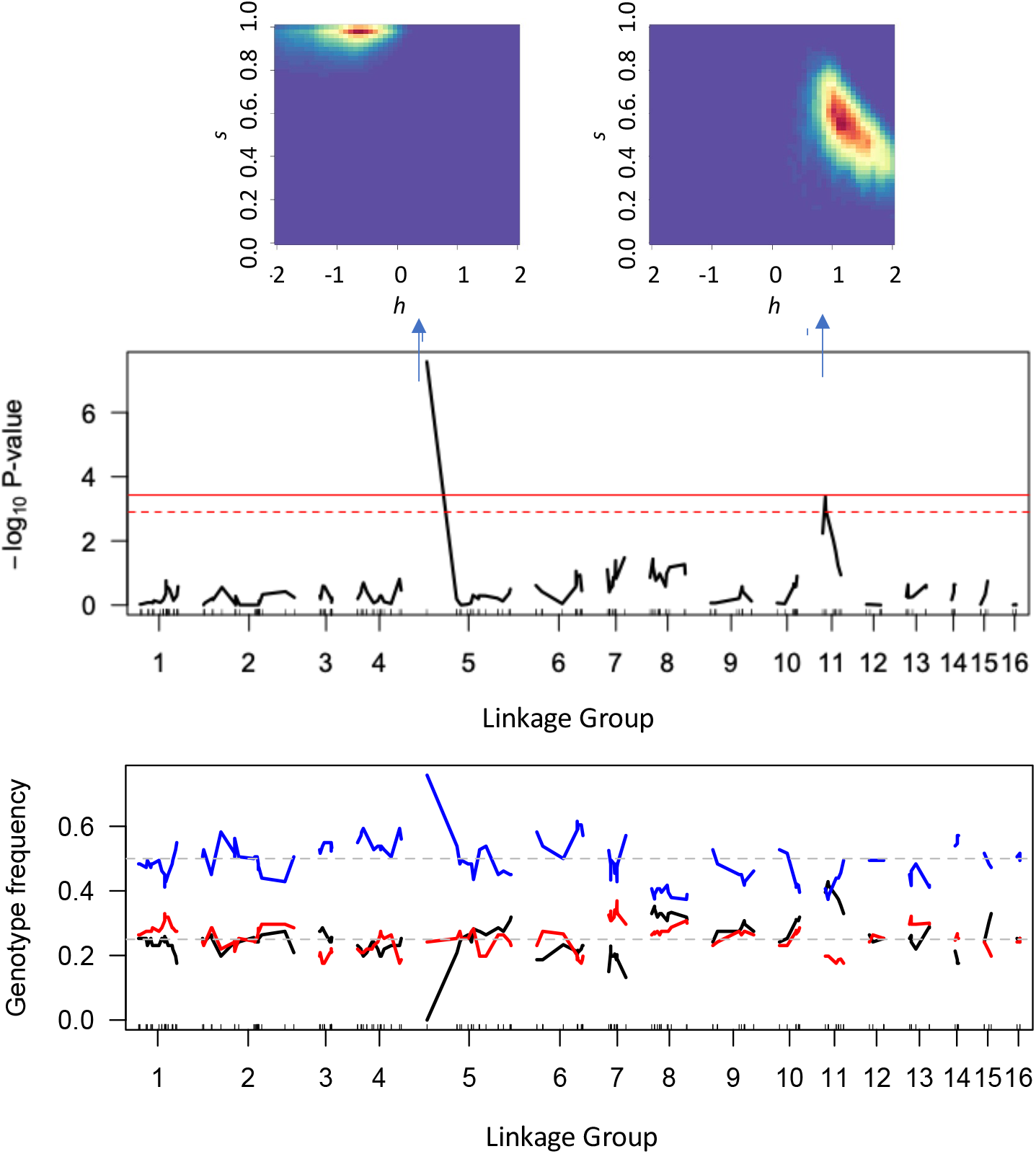
−Log_10_ significance levels from Chi-square goodness-of-fit tests to the 1:2:1 segregation ratio for SNPs in linkage groups of Family 2. Red solid and dashed lines indicate significance levels corresponding to the false discovery rate of *q* = 0.05 and 0.10, respectively. Below are the segregation ratios for the SNPs. Blue denotes heterozygote, red and black denote homozygotes. Above are heat maps of posterior probability densities for dominance (*h*) and selection (*s)* coefficients of inferred viability loci in different linkage groups (positions indicated by arrows).

### Selection and dominance of viability loci linked to SNPs

In Family 1, we detected four regions of the genome with putative viability loci (Fig. 1). In three of these regions (linkage groups 1,6, and 9), Bayesian estimates of *h* and *s* support the existence of viability loci with weakly recessive gene action and large selection coefficients (Table 1, Fig. 1). In the fourth region (linkage group 6), Bayesian estimates of *h* and *s* support the existence of a viability locus with overdominant gene action and a large selection coefficient, though it should be noted that our analysis does not permit distinguishing between overdominance and pseudo-overdominance.

**Table 1.** Marker loci that show the most significant deviations from normal segregation within separate linkage groups. Marker locus (scaffold_base pair position), linkage group (LG), and posterior probabilities of overdominant (OD), recessive (R), dominant (D), and underdominant (UD) gene actions^1^. The values for the selection coefficient (*s*) and dominance coefficient *(h)* are the Bayesian estimators (95% credible intervals).

| Family | Marker | LG | $M_1M_1$ | $M_1M_2$ | $M_2M_2$ | OD | R | D | UD | $s$ | $h$ |
| --- | --- | --- | --- | --- | --- | --- | --- | --- | --- | --- | --- |
| 1 | 61_934679 | 1 | 11 | 43 | 33 | 0.11 | 0.36 | 0.45 | 0.07 | 0.67 (0.34, 1.0) | 0.47 (-0.33, 1.14) |
| 1 | 57_279929 | 2 | 11 | 42 | 34 | 0.15 | 0.38 | 0.41 | 0.06 | 0.68 (0.28, 1.0) | 0.42 (-0.45, 1.08) |
| 1 | 28_130518 | 6 | 10 | 57 | 22 | 0.99 | 0.01 | 0 | 0 | 0.61 (0.23, 0.97) | -1.82 (-3.89, -0.16) |
| 1 | 46_1956910 | 9 | 6 | 43 | 36 | 0 | 0.51 | 0.44 | 0.01 | 0.86 (0.68, 0.98) | 0.45 (0.01, 0.79) |
| 2 | 27_153823 | 5 | 0 | 69 | 22 | 0.99 | 0.01 | 0 | 0 | 0.91 (0.63, 1.0) | -1.55 (-4.28, -0.23) |
| 2 | 8_243257 | 11 | 18 | 34 | 39 | 0 | 0.01 | 0.20 | 0.79 | 0.50 (0.20, 0.73) | 1.38 (0.74, 2.31) |
| 2 | 197_101568 | - | 0 | 85 | 0 | - | - | - | - | - | - |

<sup>1</sup> Cumulative posterior probabilities associated with overdominant (OD), recessive (R), dominant (D), and underdominant (UD) modes of gene actions were determined by summing over the posterior probabilities associated with values of $h < 0$ , $0 \leq h < 0.5$ , $0.5 \leq h < 1$ , and $h \geq 1$ , respectively.

In Family 2, we detected two regions of the genome with putative viability loci (Fig. 2) and a third that we were unable to map to any linkage group. In linkage group 5, Bayesian estimates of *h* and *s* support the existence of a viability locus with overdominant gene action and a large selection coefficient, though again it should be noted that our analysis does not permit distinguishing between overdominance and pseudo-overdominance (Table 1, Fig. 2). In linkage group 11, a viability locus with underdominant gene action and a large selection coefficient was inferred (Table 1, Fig. 2). The two genomic regions where these viability loci were detected are in separate linkage groups from those detected in Family 1 (Table S3). The segregation ratio for the unmapped viability locus suggests strong overdominance. Though it is also possible that this is an artifact arising from a locus that maps to duplicated genomic regions as discussed in the Methods section (Table 1).

### The S-locus region of *Leavenworthia alabamica* and locations of viability loci

The tightly linked S-locus genes *Lal2* and *SCRL* along with their intervening non-coding region are located on scaffold 46, at base pair positions 2,035,545 to 2,044,046. Several genomic regions on this scaffold in Family 1 showed a strong deviation from the 1:2:1 ratio (Fig. 2 and Table 2). The segregation data here support recessive gene action and strong selection *h* = 0.18 to 0.45, and *s* = 0.80 to 0.86 (Table 2). In Family 2, however, this region did not harbour any detectable viability loci.

**Table 2.** SNPs on the scaffold containing the *Leavenworthia alabamica* S-locus *(Lal2)* that show significant deviations from 1:2:1 segregation. SNP locus (scaffold_base pair position) and posterior probabilities of overdominant (OD), recessive (R), dominant (D), and underdominant (UD) gene actions^1^. The values for the selection coefficient (*s*) and dominance coefficient (*h*) are the Bayesian estimators (90% credible intervals).

| SNP | OD | R | D | UD | *s* | *h* |
| --- | --- | --- | --- | --- | --- | --- |
| 46_1,497,122 | 0.26 | 0.51 | 0.23 | 0.00 | 0.80 (0.52, 0.98) | 0.18 (-0.76, 0.71) |
| 46_1,807,312 | 0.09 | 0.52 | 0.39 | 0.00 | 0.82 (0.59, 0.98) | 0.39 (-0.18, 0.80) |
| 46_1,807,671 | 0.10 | 0.56 | 0.33 | 0.01 | 0.80 (0.53, 0.95) | 0.36 (-0.16, 0.79) |
| 46_1,956,910 | 0.04 | 0.51 | 0.44 | 0.01 | 0.86 (0.68, 0.98) | 0.45 (0.01, 0.79) |
| 46_1,957,161 | 0.09 | 0.52 | 0.38 | 0.01 | 0.85 (0.64, 0.96) | 0.38 (-0.18, 0.80) |

<sup>1</sup> Base pair position of the start of exon 1 of *LalSCRL* on scaffold 46 is 2,035,545 bp

### Statistical power to detect different modes of gene action

Statistical power to detect different modes of gene action in experiments of the type conducted here depends on the value of the recombination fraction *c*. For close linkage, e.g., *c* near 0 as is likely for most SNPs in this study (Figs. 1 and 2, Table S2), overdominant and underdominant action are easier to detect than dominant and recessive action (Fig. 3). As the recombination fraction between marker and inbreeding loci increases, statistical power to detect underdominant selection increases, whereas statistical power to detect overdominant gene action decreases (Fig. 3). For dominant and recessive gene actions, statistical power of detection is reduced with increasing *c*, but compared with overdominant and underdominant action the changes are less affected by recombination fraction. Given a sample size of 90 progeny, close linkage, and overdominant or underdominant genes actions, selection coefficients of *ca*. *s* = 0.3 at viability loci are detectable with statistical power of 80%. For dominant and recessive actions, viability loci with selection coefficients of *ca*. *s* = 0.6 are detectable with statistical power of 80%.

**Figure 3.**
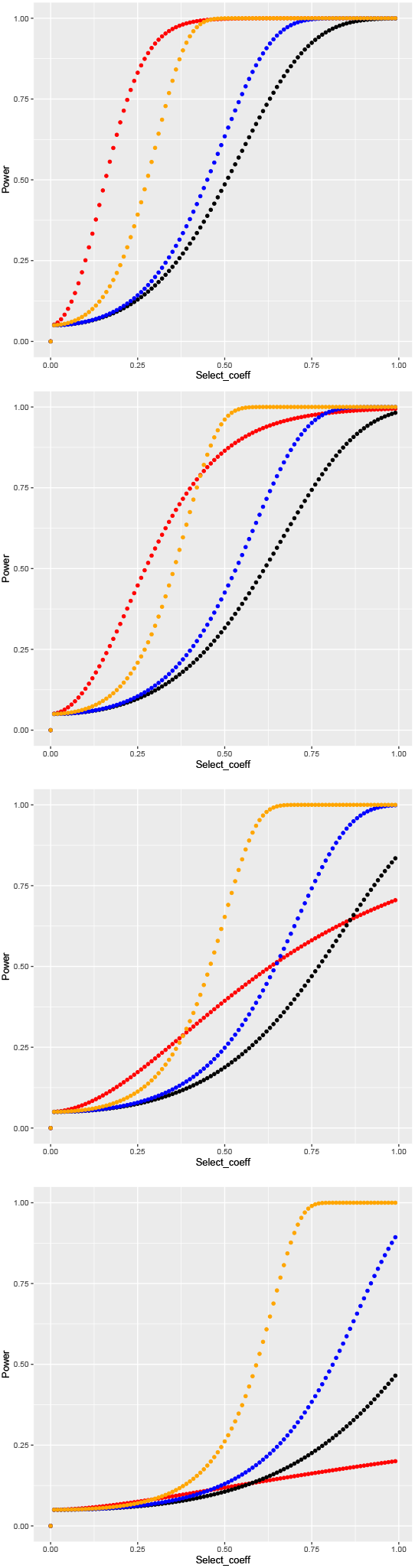
Statistical power to reject the 1:2:1 segregation ratio for different modes of action of viability loci, selection coefficients, and recombination fractions. Shown are four cases: overdominant with *h* = −3.0 (red), recessive with *h* = 0.25 (black), dominant (blue) with *h* = 0.75, and underdominant (*h* = 1.5) (orange). From top to bottom graphs, the values of the recombination fraction are *c* = 0, 0.1, 0.2, and 0.3. The graphs are based on a sample size of 90 progeny.

## Discussion

### The genetic architecture of inbreeding depression

We detected a number of significant departures from the 1:2:1 segregation ratio at SNPs located in different regions of the *Leavenworthia alabamica* genome in the progeny of parental plants from two SI populations. The segregation data were used to explore support for different values for selection and dominance coefficients associated with linked viability loci. The approach we used assumes that meiotic drive, gametic selection, and epistasis do not contribute to departures from normal segregation. The inability to distinguish the effect of these factors from viability selection is a limitation of this approach. Nevertheless, there are several reasons to believe that most departures from the 1:2:1 segregation ratio observed in this study are associated with inbreeding depression rather than with meiotic drive or gametic selection. First, where evidence of meiotic drive has been studied in detail in other species (e.g., in *Arabidopsis thaliana*), the effects were generally found in specific genomic regions, rather than genome wide as observed in the present study (Salomé *et al*., 2012; Seymour *et al*., 2019). Second, gametic selection favouring a specific allele linked to markers is not expected to produce a heterozygote excess at linked marker loci, as observed in this study for some of the marker loci in this study.

Several past studies have used marker data to explore the genetic architecture of inbreeding depression in plants. Hedrick & Muona (1990) used the approach in *Pinus sylvestris* and detected a single viability locus with a strong contribution to the fitness of progeny. In another study, using progeny from self-pollinated plants collected from a wild *Mimulus guttatus* population, Fu & Ritland (1994b) applied a graphical approach that divides the space of possible segregation ratios into different modes of gene action. They detected 24 viability loci, with the majority showing underdominance or dominance effects. Mitchell-Olds (1995), studying inbreeding depression in the progeny of parent of an *Arabidopsis thaliana* F1 hybrid between different accessions, reported a single viability locus with overdominance that mapped to chromosome 1. Remington and O’Malley (2000) examined progeny segregation ratios at AFLP loci in selfed *Pinus taeda* plants and detected both recessive and overdominant action at 19 embryonic viability loci. In the only other study to apply a Bayesian approach to estimating selection and dominance at loci associated with inbreeding depression, albeit with a different parameterisation that the one employed here, Kärkkäinen *et al*. (1999) in *Arabidopsis lyrata* ssp*. petraea* reported linkages of viability loci to seven separate allozyme loci, mostly with overdominant effects. Finally, Hedrick *et al*. (2016) examined segregation ratios for nearly 10,000 SNPs in 28 progeny from a self-pollinated plant of *Eucalytus grandis*, and found widespread heterozygote excess, consistent with overdominance, though their simulation analysis suggested that pseudo-overdominance is the more likely explanation. Overall, these empirical studies suggest that modes of gene action for linked viability loci may vary from species to species.

It is notable that a few of the above-cited studies have reported underdominant viability loci linked to markers (e.g., especially Fu & Ritland, 1994b, but also Kärkkäinen *et al*. 1999). Underdominance was also inferred at two marker-linked viability loci in the present investigation. Underdominance at viability loci cannot contribute to inbreeding depression. We note from the results of the power analyses we conducted, underdominance is detectable with smaller sample sizes compared with other modes of action that require larger sample sizes to detect selection of the same magnitude when *c* < 0.3 (Fig. 3). This could help account for the unexpected reports of underdominance in studies that have employed few markers (e.g., Fu & Ritland, 1994b).

### Overdominance or pseudo-overdominance of inbreeding depression

For a number of the marker loci showing departures from normal segregation, our analysis suggests overdominance at linked viability loci contributes to inbreeding depression. Alternatively, these results could be explained as being due to pseudo-overdominance (Ohta, 1971). Psuedo-overdominance would be detected in a genomic region by the methods used here if deleterious, recessive viability mutations are linked and in repulsion. In general, compared with recessive or partly recessive action at viability loci, there is little support from past studies for true overdominance as an underlying cause of inbreeding depression (Charlesworth and Willis 2009). Distinguishing between true overdominance and pseudo-overdominance is difficult but becomes progressively easier with multiple families. True overdominance, if present, should often be detected at the same viability loci in the progeny of multiple selfed parents, since selection there should render it frequent. In contrast, viability loci exhibiting pseudo-overdominance arising from deleterious recessive or partly recessive alleles are less likely to be shared among parents, as the genomic regions are maintained by the interaction of mutation and selection. This leaves open the possibility that both true and pseudo-overdominance are present.

### Sheltered S-linked load

An inferred viability locus (or possibly, multiple linked loci) with recessive action was detected only in one genomic region, and one family (Family 1). Interestingly, the region is within 80 kb from the site of the S-locus. These results can be interpreted as evidence that there is sheltering from selection of one (or more) recessive, deleterious mutations in the S-locus region (Table 2).

In Family 2, no such effects were detected in the S-locus region. The finding is not necessarily at odds with the evidence of sheltered load in the Family 1 data. There are likely several levels of dominance for S-locus alleles in *L. alabamica* (Busch *et al*., 2010), and in sporophytic systems such as this one, the strongest sheltering from selection is expected for those viability loci linked to S-alleles that exhibit the highest degree of dominance (Llaurens *et al*., 2009).

### Inbreeding depression, purging, and the maintenance of self-incompatibility in *Leavenworthia alabamica*

Busch (2005) found significant levels of inbreeding depression for germination and survival rate, biomass, flower number, and pollen viability in self-incompatible populations of *Leavenworthia alabamica*, and estimated that inbreeding depression for total lifetime fitness exceeds 0.5. Baldwin & Schoen (2019) studied inbreeding depression for a similar set of fitness components in *L. alabamica* and again found that inbreeding depression for lifetime fitness exceeds 0.5. They extended their study to include three consecutive generations of self-pollination and found no evidence that inbreeding depression was purged by this treatment.

Inbreeding depression is thought to be the main barrier to the evolution of self-pollination (Fisher, 1941; Lloyd 1979), and yet with some self-pollination, purging of recessive load is expected. In an earlier set of population surveys Baldwin & Schoen (2017) found that some selfing likely does occur in populations of *Leavenworthia alabamica* that have been formerly characterized at SI. Between 6% and 18% of plants exhibit “leaky self-incompatibilty” in which pollen tubes from self-pollen penetrates the styles. Such plants are presumably capable of setting seed with self-pollen. Baldwin & Schoen (2017) also showed that variation in the degree of this leakiness is heritable, and so at least one requirement for purging inbreeding depression (i.e., heritable variation for level of self-pollination) is likely present in populations of this species. This raises the question of why SI is maintained. Theory suggests that the spread of mutations that reduce SI occurs mainly when inbreeding depression loci are of major effect and recessive (Porcher & Lande, 2005; Gervais *et al*., 2014). The nearly additive viability loci detected in this study are not expected to contribute significantly to standing inbreeding depression (Charlesworth & Charlesworth, 1987). Moreover, any component of inbreeding depression due to loci with overdominance or pseudo-overdominance (as also detected in this study) should be resistant to purging. Given our findings here, we suggest that inbreeding depression in the self-incompatible populations of this species is likely due to many loci of small effect—or in other words, with effects too mild to be detected in the type of study we reort here. Inbreeding depression of this type would be expected to be resistant to purging (Charlesworth & Charlesworth, 1987) as we observed in our earlier work (Baldwin & Schoen, 2019).

## Acknowledgements

We thank Mark Romer and Claire Cooney for assistance with plant growth in the McGill Phytotron. Yong-Bi Fu provided comments on the manuscript. SJB acknowledges scholarship support from the Fonds de Recherche du Québec – Nature et technologies and support from a Discovery Grant from the Natural Sciences and Engineering Research Council of Canada to DJS

## Author Contributions

DJS and SJB conceived the study. DJS developed the Bayesian approach for characterizing the genetic architecture of viability loci underlying inbreeding depression, and the power analysis.

SJB and DJS initiated the study. Both authors analyzed and interpreted the data and wrote the manuscript. The authors are joint first authors.

## Data Availability

The *Leavenworthia alabamica* reference assembly version 2.5 is available on the CoGe platform: https://genomevolution.org/coge/GenomeInfo.pl?gid=20247. FASTQ files of GBS sequence data for parents and progeny are available on GenBank under SRA accession number PRJNA813545

## Supporting Information

| <u>File name</u> | <u>Description</u> |
| --- | --- |
| TableS1.xlsx | Table S1. Scripts and additional information used to process GBS data in STACKS |
| TableS2.xlsx | Table S2. SNPs from Families 1 and 2, their position in linkage groups and scaffolds; and segregation patterns. |
| TableS3.xlsx | Table S3. Cross reference for linkage groups in Families 1 and 2. |
| Note 1 | R code for Bayesian analysis |
| Note 2 | R code for power analysis in Fig. 3 |

## Notes

### Competing Interest Statement

The authors have declared no competing interest.

### Summary of Updates

Typos and grammatical corrections. Filtering of SNPs with high and low allele balance before data analysis.

