## Supplementary material for "The maintenance of self-incompatibility and the genetic architecture of inbreeding depression": Supplmental Note 1

```
# Power analysis for detection of departure from 1:2:1 segregation
# under varying dominance (h) and selection (s) and linkage fraction (c)
# for a viability locus linked to a marker (Fig. 4)
library(pwr)
library(ggplot2)

Power <- vector("numeric", 100)
Select_coeff <- vector("numeric", 100)
c=.3

h=-3

for (i in 1:99) {s=i/100.
Select_coeff[i] <- i/100.
# pij's are segregation proportions
p11=(((c^2)*s*(2*h-1)-2*c*s*(h-1)+(1-s))/(4-s*(2*h+1)))
p12=(2*((-c^2)*s*(2*h-1) + c*s*(2*h-1)-h*s+1)/(4-s*(2*h+1)))
p22=(((c^2)*s*(2*h-1)-2*c*s*h+1)/(4-s*(2*h+1)))

alt = (c(p11, p12, p22))
null <- c(0.25, .5, .25)
ES.w1(null,alt)
pw <- pwr.chisq.test(w=ES.w1(null,alt), N=90, df=(3-1), sig.level=0.05)
Power[i] <- pw$power
}

plod <- data.frame(cbind(Select_coeff, Power))

Power <- vector("numeric", 100)
Select_coeff <- vector("numeric", 100)
c=.3

h=0.25

for (i in 1:99) {s=i/100.
Select_coeff[i] <- i/100.
p11=(((c^2)*s*(2*h-1)-2*c*s*(h-1)+(1-s))/(4-s*(2*h+1)))
p12=(2*((-c^2)*s*(2*h-1) + c*s*(2*h-1)-h*s+1)/(4-s*(2*h+1)))
p22=(((c^2)*s*(2*h-1)-2*c*s*h+1)/(4-s*(2*h+1)))

alt = (c(p11, p12, p22))
null <- c(0.25, .5, .25)
ES.w1(null,alt)
pw <- pwr.chisq.test(w=ES.w1(null,alt), N=90, df=(3-1), sig.level=0.05)
Power[i] <- pw$power
}

plrec <- data.frame(cbind(Select_coeff, Power))
```

```


Power <- vector("numeric", 100)
Select_coeff <- vector("numeric", 100)
c=.3

h=0.75

for (i in 1:99) {s=i/100.
Select_coeff[i] <- i/100.
p11=(((c^2)*s*(2*h-1)-2*c*s*(h-1)+(1-s))/(4-s*(2*h+1)))
p12=(2*((-c^2)*s*(2*h-1) + c*s*(2*h-1)-h*s+1)/(4-s*(2*h+1)))
p22=(((c^2)*s*(2*h-1)-2*c*s*h+1)/(4-s*(2*h+1)))

alt = (c(p11, p12, p22))
null <- c(0.25, .5, .25)
ES.w1(null,alt)
pw <- pwr.chisq.test(w=ES.w1(null,alt), N=90, df=(3-1), sig.level=0.05)
Power[i] <- pw$power
}

pldm <- data.frame(cbind(Select_coeff, Power))

Power <- vector("numeric", 100)
Select_coeff <- vector("numeric", 100)
c=.3

h=1.5

for (i in 1:99) {s=i/100.
Select_coeff[i] <- i/100.
p11=(((c^2)*s*(2*h-1)-2*c*s*(h-1)+(1-s))/(4-s*(2*h+1)))
p12=(2*((-c^2)*s*(2*h-1) + c*s*(2*h-1)-h*s+1)/(4-s*(2*h+1)))
p22=(((c^2)*s*(2*h-1)-2*c*s*h+1)/(4-s*(2*h+1)))

alt = (c(p11, p12, p22))
null <- c(0.25, .5, .25)
ES.w1(null,alt)
pw <- pwr.chisq.test(w=ES.w1(null,alt), N=90, df=(3-1), sig.level=0.05)
Power[i] <- pw$power
}

plud <- data.frame(cbind(Select_coeff, Power))



Power <- vector("numeric", 100)
Select_coeff <- vector("numeric", 100)
```

```
c=.3

h=-3

for (i in 1:99) {s=i/100.
Select_coeff[i] <- i/100.
p11=(((c^2)*s*(2*h-1)-2*c*s*(h-1)+(1-s))/(4-s*(2*h+1)))
p12=(2*((-c^2)*s*(2*h-1) + c*s*(2*h-1)-h*s+1)/(4-s*(2*h+1)))
p22=(((c^2)*s*(2*h-1)-2*c*s*h+1)/(4-s*(2*h+1)))

alt = (c(p11, p12, p22))
null <- c(0.25, .5, .25)
ES.w1(null,alt)
pw <- pwr.chisq.test(w=ES.w1(null,alt), N=90, df=(3-1), sig.level=0.05)
Power[i] <- pw$power
}

plod <- data.frame(cbind(Select_coeff, Power))

Power <- vector("numeric", 100)
Select_coeff <- vector("numeric", 100)
c=.3

h=0.25

for (i in 1:99) {s=i/100.
Select_coeff[i] <- i/100.
p11=(((c^2)*s*(2*h-1)-2*c*s*(h-1)+(1-s))/(4-s*(2*h+1)))
p12=(2*((-c^2)*s*(2*h-1) + c*s*(2*h-1)-h*s+1)/(4-s*(2*h+1)))
p22=(((c^2)*s*(2*h-1)-2*c*s*h+1)/(4-s*(2*h+1)))

alt = (c(p11, p12, p22))
null <- c(0.25, .5, .25)
ES.w1(null,alt)
pw <- pwr.chisq.test(w=ES.w1(null,alt), N=90, df=(3-1), sig.level=0.05)
Power[i] <- pw$power
}

plrec <- data.frame(cbind(Select_coeff, Power))


Power <- vector("numeric", 100)
Select_coeff <- vector("numeric", 100)
c=.3

h=0.75

for (i in 1:99) {s=i/100.
```

```
Select_coeff[i] <- i/100.
p11=(((c^2)*s*(2*h-1)-2*c*s*(h-1)+(1-s))/(4-s*(2*h+1)))
p12=(2*((-c^2)*s*(2*h-1) + c*s*(2*h-1)-h*s+1)/(4-s*(2*h+1)))
p22=(((c^2)*s*(2*h-1)-2*c*s*h+1)/(4-s*(2*h+1)))

alt = (c(p11, p12, p22))
null <- c(0.25, .5, .25)
ES.w1(null,alt)
pw <- pwr.chisq.test(w=ES.w1(null,alt), N=90, df=(3-1), sig.level=0.05)
Power[i] <- pw$power
}

pldm <- data.frame(cbind(Select_coeff, Power))

Power <- vector("numeric", 100)
Select_coeff <- vector("numeric", 100)
c=.3

h=1.5

for (i in 1:99) {s=i/100.
Select_coeff[i] <- i/100.
p11=(((c^2)*s*(2*h-1)-2*c*s*(h-1)+(1-s))/(4-s*(2*h+1)))
p12=(2*((-c^2)*s*(2*h-1) + c*s*(2*h-1)-h*s+1)/(4-s*(2*h+1)))
p22=(((c^2)*s*(2*h-1)-2*c*s*h+1)/(4-s*(2*h+1)))

alt = (c(p11, p12, p22))
null <- c(0.25, .5, .25)
ES.w1(null,alt)
pw <- pwr.chisq.test(w=ES.w1(null,alt), N=90, df=(3-1), sig.level=0.05)
Power[i] <- pw$power
}

plud <- data.frame(cbind(Select_coeff, Power))


ggp <- ggplot(NULL, aes(Select_coeff,Power)) + geom_point(data = plod, col= "Red") + geom_point(data = plrec, col= "Black")+ geom_point(data = pldm, col= "Blue") + geom_point(data = plud, col= "orange")
ggp
```
