## Supplementary material for "The maintenance of self-incompatibility and the genetic architecture of inbreeding depression": Supplmental Note 2

```
#Bayesian analysis of marker segregation ratios for linked viability
#locus; support for different values of h and s (Fig 3, Tables 1 and 2)

rm(list=ls())
setwd("/Users/schoen/Desktop")
library(MASS)
library(RColorBrewer)
library(writexl)
# set prior for selection coefficient (s)
priors <- function(s) {
  if ((s < 0) || (s > 1)) { # || here means "or"
    return(0)
  } else {
    return(1)
  }
}

# set prior for domiance coefficient (h)
priorh <- function(h) {
  if ((h < -100) || (h > 100)) { # || here means "or"
    return(0)
  } else {
    return(1)
  }
}

# set prior for recombination fraction (c)
priorc <- function(c) {
  if ((c < 0) || (c > 0.20)) { # || here means "or"
    return(0)
  } else {
    return(1)
  }
}

likelihood <- function(c,s,h, naa, nAa, nAA) {

    return((((c^2)*s*(2*h-1)-2*c*s*(h-1)+(1-s))/(4-s*(2*h+1)))^naa * (2*((-c^2)*s*(2*h-1) + c*s*(2*h-1)-h*s+1)/(4-s*(2*h+1)))^nAa * (((c^2)*s*(2*h-1)-2*c*s*h+1)/(4-s*(2*h+1)))^nAA)

           }

shsampler <- function(naa, nAa, nAA, niter,cstartval,
                      sstartval, hstartval,cproposalsd,
                      sproposalsd, hproposalsd) {
    s <- rep(0,niter)
    h <- rep(0,niter)
```

```
    c <- rep(0,niter)
    s[1] <- sstartval
    h[1] <- hstartval
    c[1] <- cstartval
    for (i in 2:niter) {
        currents <- s[i - 1]
        currenth <- h[i - 1]
        currentc <- c[i - 1]


        news <- currents + rnorm(1, 0, sproposalsd)


        A <- priors(news) * likelihood(currentc,news,currenth,naa,nAa,nAA) /
          (priors(currents) * likelihood(currentc,currents,currenth,naa,nAa,nAA))



          if (runif(1) < A) {
          s[i] <- news
        } else {
          s[i] <- currents
        }



        newh <- currenth + rnorm(1, 0, hproposalsd)

        B <- priorh(newh) * likelihood(currentc,s[i],newh,naa,nAa,nAA) /
          (priorh(currenth) * likelihood(currentc,s[i],currenth,naa,nAa,nAA))
          if (runif(1) < B) {
          h[i] <- newh
        } else {
          h[i] <- currenth
        }

        newc <- currentc + rnorm(1, 0, cproposalsd)

        C <- priorc(newc) * likelihood(newc,s[i],h[i],naa,nAa,nAA) /
          (priorc(currentc) * likelihood(currentc,s[i],h[i],naa,nAa,nAA))
        if (runif(1) < C) {
          c[i] <- newc
        } else {
          c[i] <- currentc
        }
```

```
    }
    return(list(s = s,h = h,c = c)) # return a "list" with three elements named s, h, and c
}



x <- seq(0, 1, length = 3000000)

# enter segregation data on next line, e.g. 6, 43, 36 for M1/M1, M1/M2, and M2/M2 genotype numbers where relative fitnesses are (1-s), (1-hs) and 1
# followed by number of iterations (e.g., 6,000,000), followed parameters used to control the Metropolis-Hastings sampler.
z <- shsampler(10, 57, 22, 3000000, .05, .5, 0, .01,.01,.01)

# draw histograms (adjust xlim and ylim according to expected results). Histograms based on last 500,000 iterations after the burnin at 5,500,000)
jpeg('plot_6_43_36.jpg')
par(mfrow = c(1,2))
hist(z$s[2500000:3000000], prob = T, xlim = c(0, 1), col = "dark grey", ylim = c(0,2), main = "s sample", xlab = "posterior estimate", freq=FALSE)
lines(density(z$s[2500000:3000000])) # density plotlwd = 2, thickness of linecol = "chocolate3")
hist(z$h[2500000:30000000], prob = T, xlim = c(-1, 2), col = "dark grey", ylim = c(0,1),  main = "h sample", xlab = "posterior estimate")
lines(density(z$h[2500000:3000000])) # density plotlwd = 2, thickness of linecol = "chocolate3")
dev.off()

# write last 500,000 values to Excel frequency table.  The results in these files were used to estimate values in Table 1 and Table 2.

hist_info <- hist(z$h[2500000:3000000], prob = T, breaks=seq(-3,3,.01), col = "dark grey", ylim = c(0, 10),  main = "h sample", xlab = "posterior estimate")
hist_info$mids
hist_info$counts
hdist <- data.frame(rbind(hist_info$mids,hist_info$counts))
hdist <- t(hdist)
hdist <- data.frame(hdist)
write_xlsx(hdist ,path =  "hdist_6_43_36.xlsx",col_names = TRUE,format_headers = TRUE,use_zip64 = FALSE)

hist_info <- hist(z$s[2500000:3000000], prob = T, breaks=seq(-.01,1,.01), col = "dark grey", ylim = c(0, 10),  main = "s sample", xlab = "posterior estimate")
hist_info$mids
```

```
hist_info$counts
sdist <- data.frame(rbind(hist_info$mids,hist_info$counts))
sdist <- t(sdist)
sdist <- data.frame(sdist)
write_xlsx(sdist ,path =  "sdist_6_43_36.xlsx",col_names = TRUE,format_headers = TRUE,use_zip64 = FALSE)

x=z$h[2500000:3000000]
#range(x)
y=z$s[2500000:3000000]
#range(y)
df=data.frame(x,y)

# draw "heat map" of probability densities for h and s
jpeg('plot3D_6_43_36.jpg')
par(mfrow = c(1,1))
rf <- colorRampPalette(rev(brewer.pal(11,'Spectral')))
r <- rf(63)
k <- kde2d(df$x, df$y, n=50,lims = c(-5,2,0,1))
image(k, col=r)
dev.off()
```
